# Chromatin architecture reorganisation during neuronal cell differentiation in Drosophila genome

**DOI:** 10.1101/395822

**Authors:** Keerthi T Chathoth, Nicolae Radu Zabet

## Abstract

Compartmentalisation of the genome as topologically associating domains (TADs) may have regulatory role in development and cellular functioning, but the, mechanism involved in TAD establishment is still unclear. Here, we present the first high-resolution contact map of *Drosophila melanogaster* neuronal cells (BG3) and identified different classes of TADs by comparing this to genome organisation in embryonic cells (Kc167). We find new rearrangements during differentiation in neuronal cells reflected as enhanced long-range interactions, which is supported by pronounced enrichment of CTCF at cell type specific borders. Furthermore, we show the presence of strong divergent transcription corroborated with RNA Polymerase II occupancy and increased DNA accessibility at the TAD borders. Interestingly, TAD borders that are specific to neuronal cells are enriched in enhancers controlled by neuronal specific transcription factors. Our results suggest that TADs are dynamic across developmental stages and reflect the interplay between insulators, transcriptional states and enhancer activities.

## INTRODUCTION

Chromosome conformation capture Hi-C technique has paved way to dissect the compartmental organisation of genome in various cell types (Dekker et al. 2002; Lieberman-Aiden et al. 2009; Dixon et al. 2012; Nora et al. 2012; Dixon et al. 2015; Flyamer et al. 2017). Further advancement to high-resolution methodologies such as *Insitu* Hi-C has enabled to obtain much more refined 3D organisation of the genome from megabase scale compartments to subkilobase resolution (Rao et al. 2014; Cubeñas-Potts et al. 2017). Topologically associating domain has been regarded as an important basic unit of chromosome organisation. They are believed to be evolutionarily conserved and appear as invariant across different organisms and cell types (Rao et al. 2014; Vietri Rudan et al. 2015; Dixon et al. 2015).

Most of the self-interacting regions observed within and between TADs are shown to arise from promoter enhancer interactions (Noordermeer et al. 2014; Cubeñas-Potts et al. 2017). Such dynamic regulation of long-range contacts, required for cell differentiation, is thought to occur within TADs. Similarly, the establishment of enhancer promoter loops was shown tightly coupled to activation of poised enhancers as well as to gene expression (Freire-Pritchett et al. 2017). These internal interactions within TADs appear to change during development (Dixon et al. 2015) and under heat shock (Li et al. 2015), indicating a functional role for TADs. Although the functional importance of TADs was shown previously (Lupianez et al. 2015), the stability and establishment of borders is not yet fully understood.

TADs have been correlated with various factors. They are reported to be regions lacking active chromatin marks and separated by inactive marks (Ulianov et al. 2016; El-Sharnouby et al. 2017) and, at borders, they are shown to be enriched with house-keeping and development enhancers (Cubeñas-Potts et al. 2017). It is also shown to coincide with long-range gene regulatory modules such as genomic regulatory blocks (Harmston et al. 2017). Architectural proteins are considered as another factor that is believed to play a significant role in demarcating the TAD boundaries and their enrichment has been correlated with border strength (Van Bortle et al. 2014; Stadler et al. 2017). CTCF and Cohesion are the main architectural proteins that occupy mammalian TAD borders and their absence seems to disrupt TADs architecture unevenly, suggesting the presence of different types of borders (Schwarzer et al. 2017; Zuin et al. 2014; Nora et al. 2017). In contrast, TAD borders in *Drosophila melanogaster* are occupied by a large set of insulator proteins including CTCF, BEAF-32, Chromator, CP190 etc. (Van Bortle et al. 2014; Stadler et al. 2017). Recently, transcription is emerging as another major driver of TAD formation (Li et al. 2015; Rowley et al. 2017). Interestingly, a recent study showed that TADs appear together with transcription activation in the zygote, but blocking transcription elongation does not seem to affect TADs (Hug et al. 2017). Also synthetic induction of transcription using CRISPR/Cas9 system in mouse neuronal progenitor cells does not induce TAD boundary formation (Bonev et al. 2017).

We present here the high-resolution *insitu* Hi-C carried out in Drosophila neuronal cells that enabled accurate demarcation of TAD borders. Detailed analysis revealed different classes of TADs that are distinct based on the conservation of borders during differentiation. We also find more long-range interactions in neuronal cells than in embryonic cells and this is supported by the enrichment of insulator proteins that are involved in long-range interactions such as CTCF and Chromator. Importantly, we observed a strong enrichment for divergent transcription and RNA Polymerase II occupancy over the TAD borders. Altogether, the data presented here provide new insights into the cell specific borders that are gained during differentiation and also the interplay between divergent transcription and insulator proteins on TAD border formation in *Drosophila melanogaster.*

## RESULTS

### Characterisation of Topologically associated domains based on border conservation in Drosophila

The topologically associated domain has been analysed in *Drosophila* previously (Sexton et al. 2012; Hou et al. 2012; Li et al. 2015; Ulianov et al. 2016; Cubeñas-Potts et al. 2017; Hug et al. 2017; Eagen et al. 2017; Rowley et al. 2017) using both low and high-resolution approaches. All the previous *Insitu* Hi-C studies (generating subkilobase resolution contact maps) were conducted in embryonic cell lines (Kc167 and S2) or whole embryos (Cubeñas-Potts et al. 2017; Hug et al. 2017; Eagen et al. 2017; Rowley et al. 2017). Here we generated a high-resolution chromatin map of neuronal derived cells in *Drosophila*, by performing *Insitu* Hi-C in BG3 cells (Figure 1A). To generate this map we used a four base cutter (DpnII), which resulted in an average distance between restriction sites of approximately 500 bp (see *Methods,* Figure S1A-B). To understand the characteristics of chromatin organisation in neuronal cells, we compared *Insitu* Hi-C in BG3 cells with previously published embryo-derived cells (Kc167); see Figure 1B-C. As a result, we identified 1900 TADs in BG3 cells and 2252 TADs in Kc167 cells, which is in agreement with other studies (Cubeñas-Potts et al. 2017). This is almost four times more TADs in our Hi-C map of BG3 cells compared to previous reports from a low resolution map (Ulianov et al. 2016). Additionally, when compared to Kc167 cells, BG3 exhibited substantially higher long-range contacts and fewer short-range contacts, (Figure 1B-C), which suggests a dynamic change in chromatin and gene regulation upon differentiation.

**Figure 1.**
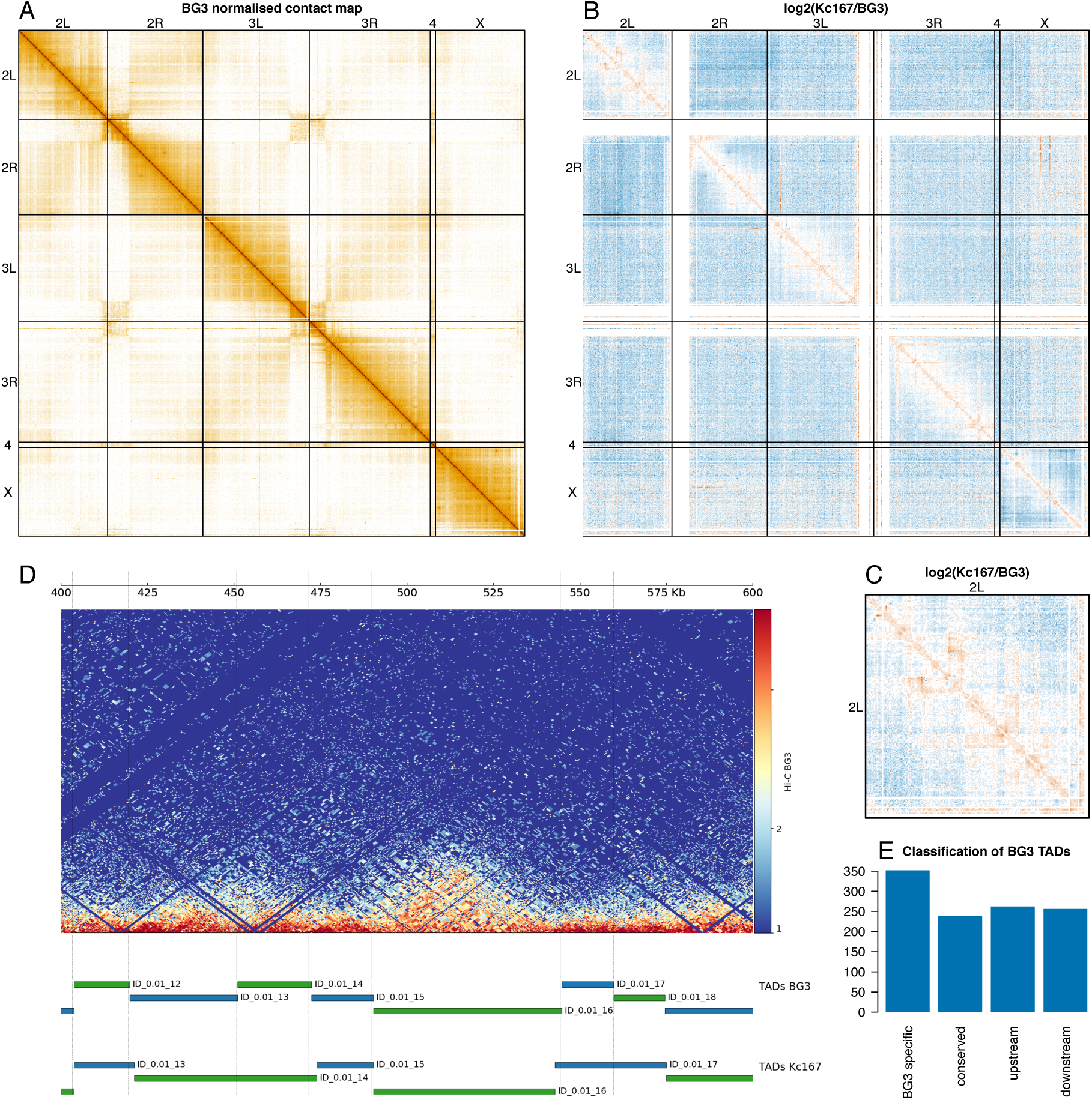
A high-resolution map of *Drosophila* BG3 cells. (A) Genome-wide normalised contact map of Drosophila BG3 cell line at 100 Kb resolution. Each element in the matrix represents the normalised number of contacts between the two corresponding bins. (B) Log2 ratio between the normalised number of contacts in BG3 cells and Kc167 cells as indicated. (C) The log2 ratio between the normalised number of contacts in BG3 cells and Kc167 cells on chromosome 2L.(D) Triangle view of the normalised contact map in BG3 cells at 2L:400,000..600,000 locus. TADs called in BG3 and Kc167 cells are displayed below for this locus as indicated. (E) Classification of TADs in BG3 cells as described in the main text: BG3 specific TADs (with both borders only present in BG3 cells), conserved TADs (with both borders located at same position in both Kc167 and BG3), upstream conserved (only the upstream border is located at same position in both cells) and downstream conserved (only the downstream border is located at same position in both cells). TADs where the borders are slightly shifted within 2 Kb (classified as fuzzy TADs) were not considered in the downstream analysis.

During TAD border analysis, one third of the borders appeared to be identical between the cell types, while the rest were positioned in varying distances (Figure S1C-D). Based on the conservation of the TAD borders in both the cell types, we classified TADs as: (*i*) BG3 specific TADs (TADs of BG3 where both borders are at least 2Kb apart from the closest borders of Kc167), (*ii*) conserved TADs (TADs with both borders conserved at the exact position in both Kc167 and BG3), (*iii*) upstream conserved TADs (BG3 TADs with only the upstream border conserved at the exact position as in Kc167 and the other border further than 2 Kb from the border in Kc167 cells) and (i*v*) downstream conserved TADs (TADs with only the downstream border conserved at the exact position as in Kc167 and the other border further than 2 Kb from the border in Kc167 cells) (Figure 1D-E). The rest of the TADs in BG3 cells have borders that are not at the exact position, but are within 2 Kb of the corresponding border in Kc167 cells. To identify the possible strong determinants for maintaining and establishing TAD boundaries, we focused on the four classes identified in Figure 1E for our downstream analysis.

### TAD borders formation is driven by divergent transcription and Polymerase occupancy

Previous work showed that TAD borders display high levels of DNA accessibility and expression (Sexton et al. 2012; Li et al. 2015), but the role of active transcription as a determinant of TAD boundaries is still disputed. The expression of genes was shown to be a major predictor of TAD boundaries (Rowley et al. 2017), whereas, blocking transcription with alpha-amanitin-treatment does not remove TAD borders (Hug et al. 2017) or synthetic induction of genes with CRISPR/Cas9 system does not lead to the appearance of TAD borders (Bonev et al. 2017). To analyse the transcriptional status between different cell types, we first compared the DNA accessibility and Pol II occupancy across different classes of TAD borders. Interestingly, DNA accessibility and Pol II occupancy was enriched at conserved borders and was comparatively high in BG3 cells at the gained BG3 specific borders (Figure 2A-H).

**Figure 2.**
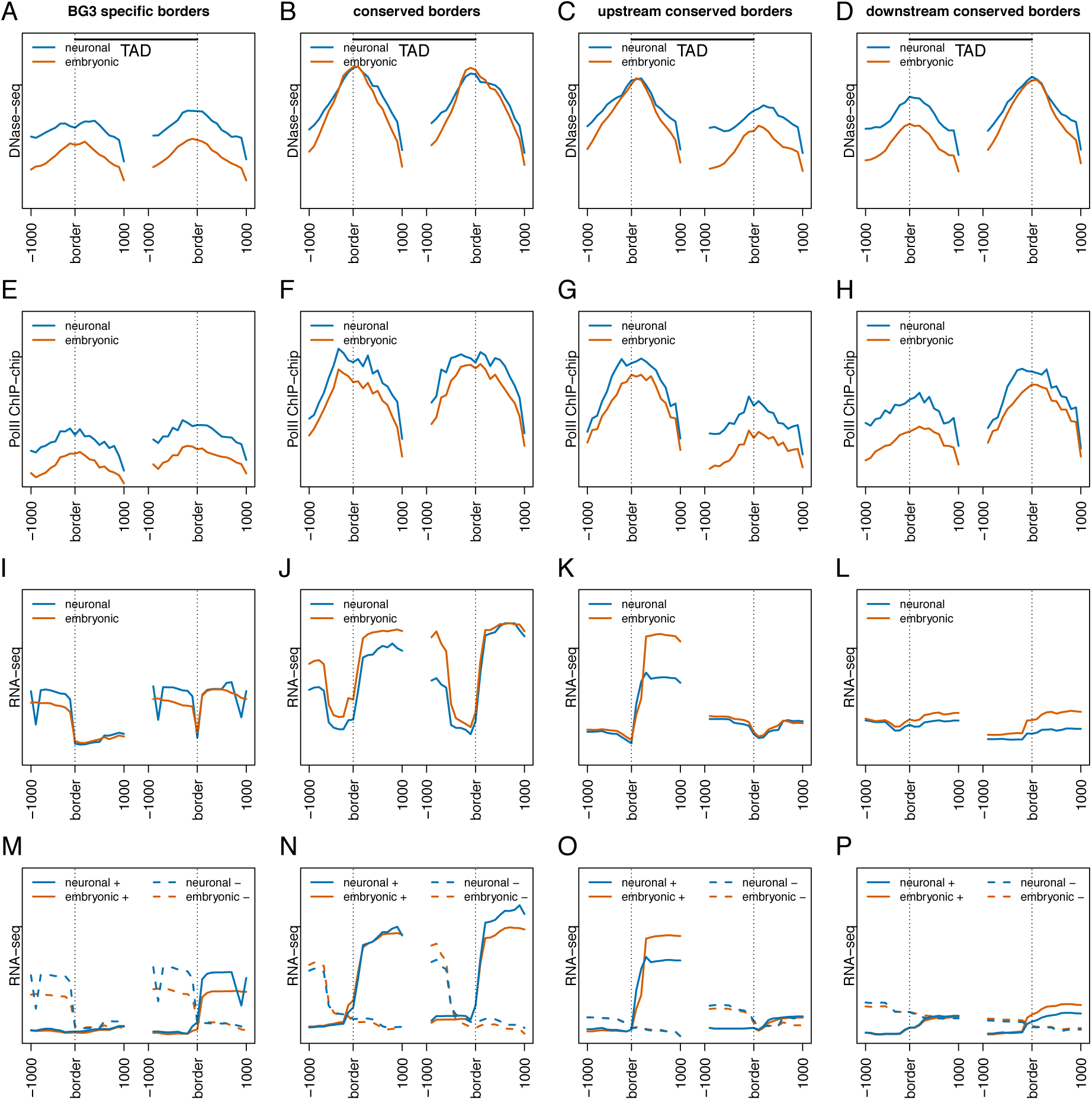
Divergent transcription and polymerase occupancy correlates with appearance of TAD borders. (A-D) DNaseI-seq signal at borders of four different TAD classes (BG3 specific, conserved, upstream conserved and downstream conserved) as indicated. Conserved borders display similar high levels of DNA accessibility, while BG3 specific TAD borders gain DNA accessibility in BG3 cells compared to Kc167 cells. Red line represents data from embryonic cells (Kc167) and blue from neuronal derived cells (BG3). Average profile has been plotted considering 1 Kb around each border. (E-H) Average PolII ChIP-chip signal (log2 ChIP/input) at borders of the four different TAD classes as indicated. (I-L) Average RNA-seq levels at borders of the four different TAD classes as indicated. (M-P) Strand specific average RNA-seq signal at borders of the four different TAD classes showing strong divergent transcription at TAD borders. Solid lines represent the positive strand and dashed lines the negative strand. The expression levels of genes in both cell types were considered and mapped back the level of expression on corresponding strands.

Looking into the PolyA RNA in both embryonic and neuronal cells we found that despite the increased presence of Pol II, total RNA expression was enriched only at the conserved borders and not much at specific borders in both cell lines (Figure 2I-L). The little difference between the two cell types indicate that transcription alone cannot explain the appearance of TAD borders in BG3 cells, which is in agreement with (Hug et al. 2017; Bonev et al. 2017). As Pol II occupancy didn’t seem to completely explain the total RNA expression, we next analysed the data in a strand specific manner. Intriguingly, a strong divergent transcription was noticed at the conserved and BG3 specific borders, which corroborates with the enriched Pol II signal (Figure 2M-P). To check if this phenomenon is encoded in the DNA sequence, we investigated the number of annotated genes at TAD borders and observed that the divergent transcription is supported by the number of genes present at the TAD borders (Figure S2A-B). However, one exception was the right arm of the upstream border of BG3 specific TADs (Figure 2M). To account for this, we reasoned that this part of divergent transcription is contributed by ncRNAs, as the number of encoded ncRNAs appears to be highly enriched on the positive strand at these loci (Figure S2C-D). To elucidate this, we then examined the nascent RNA in both cell lines that would represent all RNA species (Figure S2E-H). Strikingly, we observed the appearance of divergent transcription at BG3 specific borders in BG3 cells (Figure S2G-H). This indicates a potential role for ncRNA in sustaining divergent transcription and, consequently, in the formation or maintenance of TAD borders.

To further validate the different aspects of divergent transcription we looked at the ratio between sense and antisense transcription. There is more antisense transcription upstream of TAD borders and more sense transcription downstream of TAD borders (Figure S3A-D). Finally, using the definition of divergent transcription as less than three times more nascent transcription on one strand compared to the other strand in 500 bp windows (Jin et al. 2017), we observed that majority of TAD borders display bidirectional transcription (Figure 3). Apparently, the percentage of borders with bidirectional transcription increases for slightly more relaxed threshold values (e.g. four times more transcription on one strand) and is higher at BG3 specific borders in BG3 cells compared to Kc167 cells (Figure 3A-D). Altogether, our data is showing that increased Pol II occupancy and divergent transcription could potentially drive TAD borders formation.

**Figure 3.**
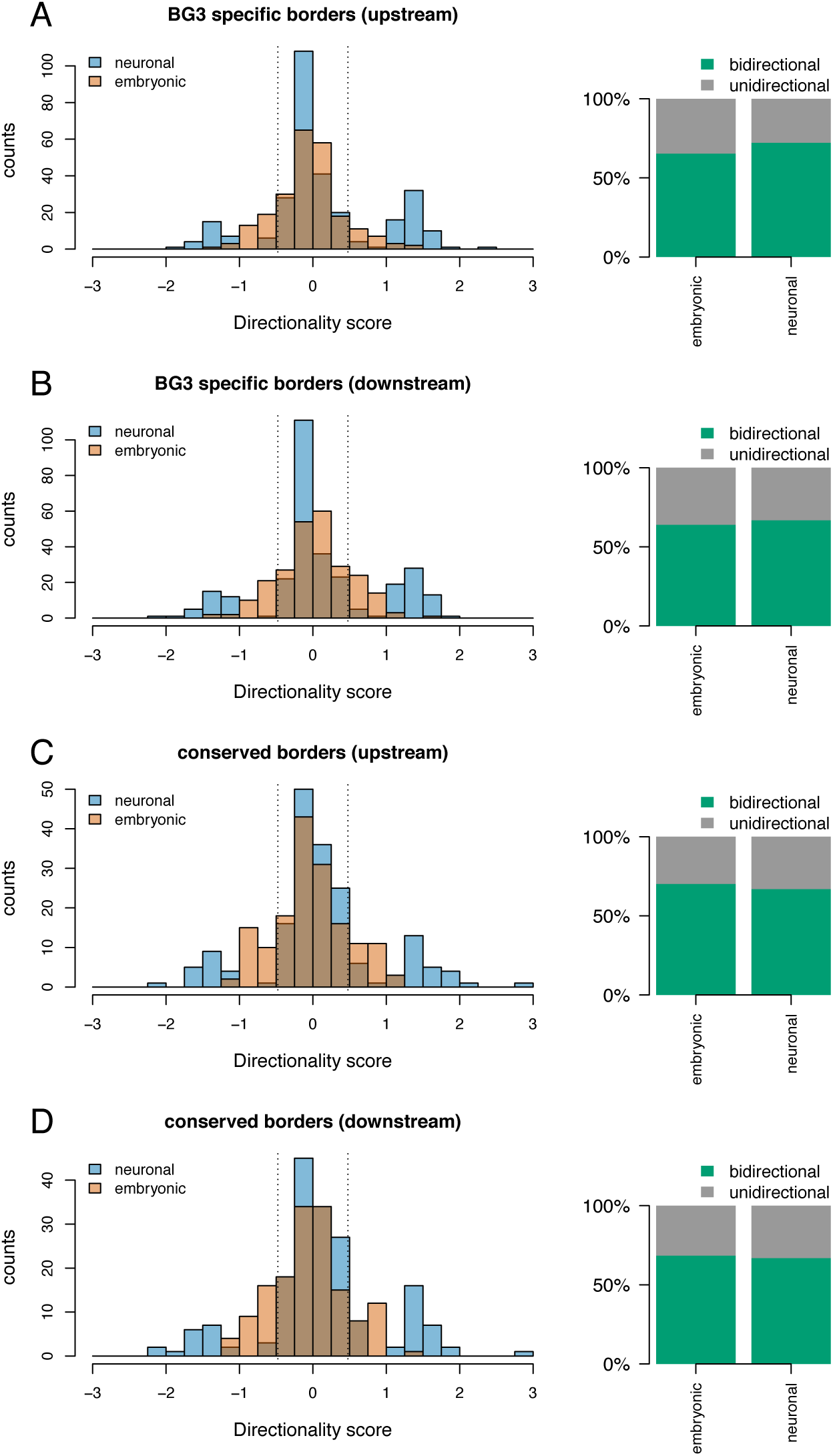
Bidirectional transcription at TAD borders. Histograms representing the directionality score computed as log10 of the ratio between nascent RNA levels in 500 bp on the positive strand downstream of the border and on the negative strand upstream of the border. 500 bp bins that were 500 bp away was considered from the border in both directions. Barplot represents the percentage of TAD borders where the directionality score was lower than 0.47 (dotted lines on the histogram, representing less than three times more transcription on one strand) and classified this as bidirectional borders. Majority of TAD borders display bidirectionality as shown. (A) upstream border of BG3 specific TADs. (B) downstream border of BG3 specific TADs. (C) upstream border of conserved TADs. (D) downstream border of conserved TADs.

### Architectural proteins are differentially occupied over borders across cell lines with CTCF being highly enriched at BG3 borders

Drosophila displays a large repertoire of architectural proteins, which are bound to TAD boundaries (Van Bortle et al. 2014). BEAF-32, CP190 and Chromator are involved in long-range interactions (Vogelmann et al. 2014) and several recent studies also found they are the most enriched proteins at TAD border in Drosophila (Cubeñas-Potts et al. 2017; Hug et al. 2017; El-Sharnouby et al. 2017; Ramirez et al. 2017). We also observed that the three insulator proteins were enriched especially at the conserved borders (Figure 4). Comparing the two cell lines, BEAF-32 and CP190 is present more at the borders of embryonic cells than in neuronal cells (Figure 4A-H); whereas, Chromator signal is high at TAD borders of neuronal versus embryonic cell lines (Figure 4 I-L).

**Figure 4.**
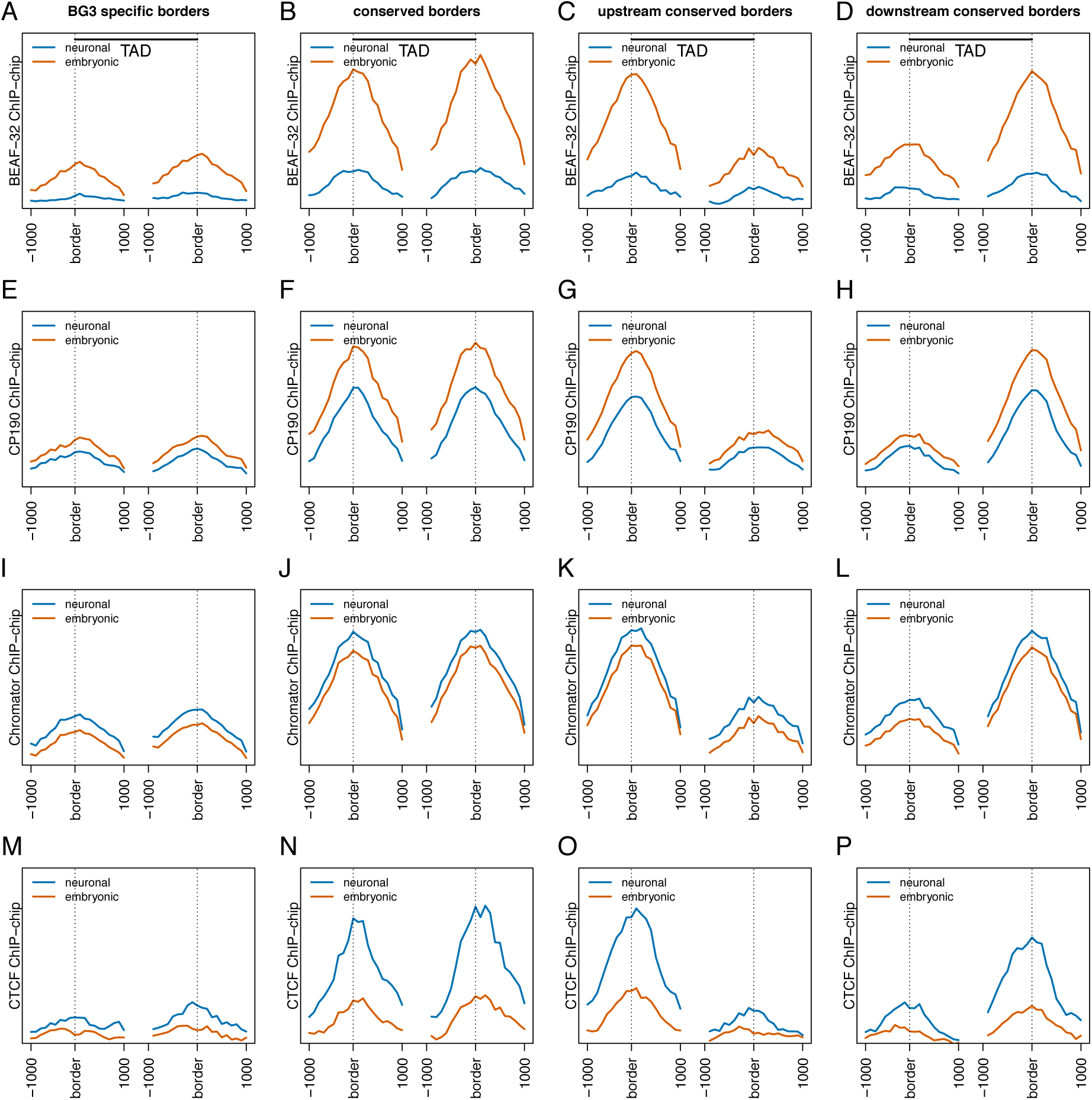
Architectural proteins differentially occupy TAD borders in a cell specific manner. (A-D) Average BEAF-32 ChIP-chip signal (log2 ChIP/input) at borders of four different TAD classes (BG3 specific, conserved, upstream conserved and downstream conserved). Red line represents data from embryonic cells and blue from neuronal derived cells. As before 1Kb around each border was considered while plotting the average profile. (E-H) Average CP190 ChIP-chip signal (log2 ChIP/input) at borders of the four different TAD classes. (I-L) Average Chromator ChIP-chip signal (log2 ChIP/input) at borders of the four different TAD classes. (M-P) Average CTCF ChIP-chip signal (log2 ChIP/input) at borders of the four different TAD classes.

In contrast to mammalian systems, previous studies reported reduced amounts of CTCF binding at TAD borders in Drosophila. However, those studies were performed either in embryonic cells (Ramirez et al. 2017) or with a low-resolution Hi-C data in differentiated cells (Ulianov et al. 2016). Strikingly, we found that CTCF binds more at TAD borders than previously reported and this increase in binding is substantially high in BG3 cells and also observed at BG3 specific TAD borders (Figure 4M-P). CTCF was shown to promote long-range interactions in Drosophila (Li et al. 2013) as well as regulate developmentally stable interactions (Phillips-Cremins et al. 2013). Given that long-range contacts are more prevalent in neuronal cells (Bonev et al. 2017), increased presence of CTCF and Chromator in Drosophila may mediate such interactions either independently or in combination with other architectural proteins.

We also examined other architectural proteins such as Su(Hw) and GAF (Figure S4). GAF profile was similar to Chromator across the border and between cell types (Figure S4A-D), while, consistent with previous reports, Su(Hw) is less enriched at TAD borders (Ulianov et al. 2016; Ramirez et al. 2017) (Figure S4E-H). The differential enrichment of certain architectural proteins may indicate their importance in TAD border establishment during different developmental stages, potentially leading to the formation and perpetuation of cell specific chromatin organisation states.

### Chromatin marks display consistent patterns at TAD borders

Several chromatin factors can work together and can act as identity for the underlying features of genes. Previous studies reported the presence of active marks (H3K27ac, H3K4me1 and H3K4me3) at the TAD borders and the presence of repressive marks inside TADs (H3K27me3) (Ulianov et al. 2016; El-Sharnouby et al. 2017; Cubeñas-Potts et al. 2017). H3K27ac is a mark enriched at active enhancers and there is an increased signal for H3K27ac in BG3 cells at specific TAD borders (Figure S5A-D). However, despite the increase in H3K27ac, the expression was low as previously shown in Figure 2, so we looked at other marks and found that there is also an increase of H3K27me3 repressive mark at the BG3 specific borders (Figure S5E-H). The presence of active and repressive marks may indicate that BG3 specific TAD borders might be in competent chromatin state (enhancer regions that contain both repressive and activating marks) (Skalska et al. 2015).

Conserved TAD borders are depleted in H3K4me1 and enriched in H3K4me3, suggesting that they are associated with actively transcribed genes (or house-keeping genes) (Skalska et al. 2015) (Figure S5E-H). In contrast, BG3 specific TAD borders display higher levels of H3K4me1 and lower levels of H3K4me3, indicating that these borders are enriched in primed enhancers (or developmental enhancers) (Skalska et al. 2015) (Figure S5). Above results suggests that while conserved TAD borders are enriched in constitutively expressed genes, the BG3 specific TAD borders could be enriched in BG3 specific enhancers.

### Enhancers are pronounced at the borders during differentiation in Drosophila

Enhancer promoter (E-P) looping is one of the contributing factors to gene regulation and is considered as one of the factors that possibly drive TAD formation (Bonev and Cavalli 2016). These E-P interactions are subjected to change during differentiation depending on the transcriptional requirement of the cells (Ghavi-Helm et al. 2014). TAD borders were classified as either being established by housekeeping genes or by developmental enhancers, in embryo derived cells (Kc167) (Cubeñas-Potts et al. 2017). Here, we could provide further evidence for this classification of TAD borders by investigating if the TAD borders that are specific to BG3 cells are enriched in neuronal associated enhancers, while conserved TAD borders between two cells are not. We used the STARR-seq data from (Arnold et al. 2013; Yanez-Cuna et al. 2014) and categorised the enhancers as BG3 specific, S2 specific and common enhancers (Figure 5A). As a result, there seem to be more BG3 specific TAD borders that contain BG3 specific enhancers than BG3 specific TAD borders that contain common or S2 enhancers, while at conserved TAD borders the number of TAD borders containing BG3 or S2 specific enhancers were similar (Figure 5B). This supports the model that enhancer promoter looping may be one of the underlying factor for BG3 specific TAD border formation.

**Figure 5.**
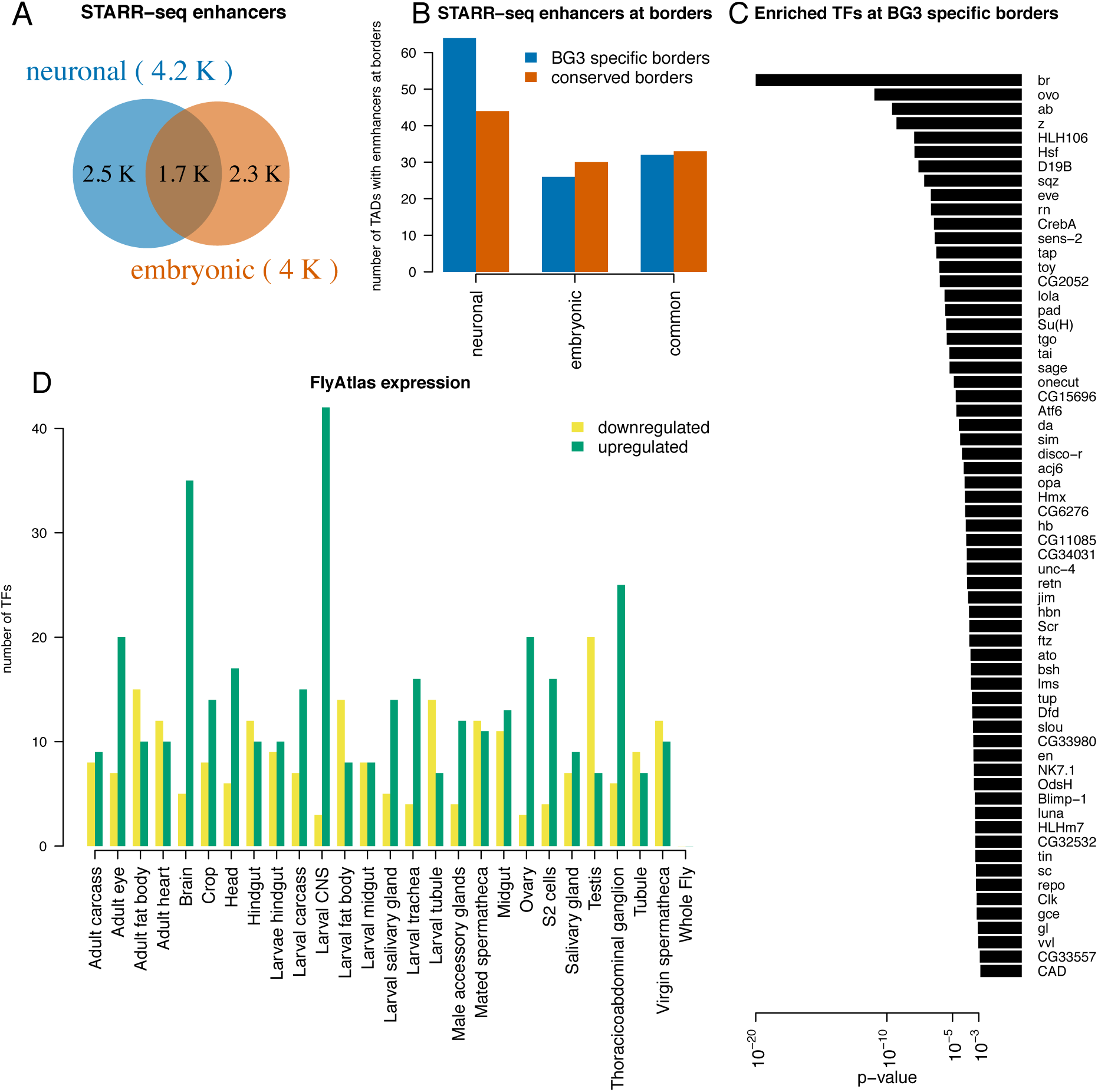
Cell specific enhancers correlate with cell specific TAD borders. (A) Venn diagram representing the number of enhancers in neuronal and embryonic Drosophila cells as listed in STARR-seq. Enhancers were classified as cell specific if they were annotated only in one cell type or common if they were annotated in both cell types. (B) The number of conserved (red) or BG3 specific (blue) TAD borders that overlap with a cell specific or common enhancer. Barplots showing more BG3 specific borders with neuronal enhancers than with common or embryonic enhancers unlike conserved TAD borders. (C) List of 63 TFs with enriched motifs at BG3 specific borders and the associated p-value (see *Methods*). (D) Expression of the 63 TFs in different Drosophila tissues/cells from FlyAtlas dataset. Green represents upregulated genes and yellow downregulated genes as indicated. The most upregulated TFs are specifically expressed in larval central nervous system (from where BG3 cells are derived) and in brain.

Nevertheless, despite this increase of neuronal specific enhancers at BG3 specific borders, a large proportion of BG3 borders were still devoid of enhancers. This was puzzling since we observed active transcription and enhancer marks at these TAD borders (Figures 2 and S5). To explain this, we used a complementary approach where we identified a list of 63 TFs that have their binding motifs enriched at BG3 specific TAD borders (see *Methods*) (Figure 5C). Using FlyAtlas dataset (Chintapalli et al. 2007), we checked if these TFs are expressed and specific to any particular tissue/cell. Notably, majority of the 63 TFs that have motifs enriched at BG3 specific TAD borders are specifically expressed in larval central nervous system (from where BG3 cells were derived) and brain (Figure 5D). This provides additional evidence that BG3 specific enhancers contribute to the formation of BG3 specific TAD borders

### BG3 exhibits more long-range contacts and Kc167 shows more short-range interaction

It was reported that *Drosophila melanogaster* displays significantly fewer chromatin loops than in mammals (Eagen et al. 2017). Our first inspection of the Hi-C map (Figure 1B) indicated that BG3 cells would have more long-range contacts, while Kc167 would display more short-range contacts. To analyse long and short-range interactions, we identified all enriched contacts in the Hi-C dataset in BG3 and Kc167 cells. As expected, we observed more enriched contacts in Kc167 cells at distances between 10 Kb and 1 Mb, and more contacts in BG3 cells for distances longer than 1 Mb (Figure 6A). Most of the TADs detected in both cells are smaller than 1Mb, indicating that BG3 cells display more contacts between TADs, while Kc167 cells display more contacts within the TADs.

**Figure 6.**
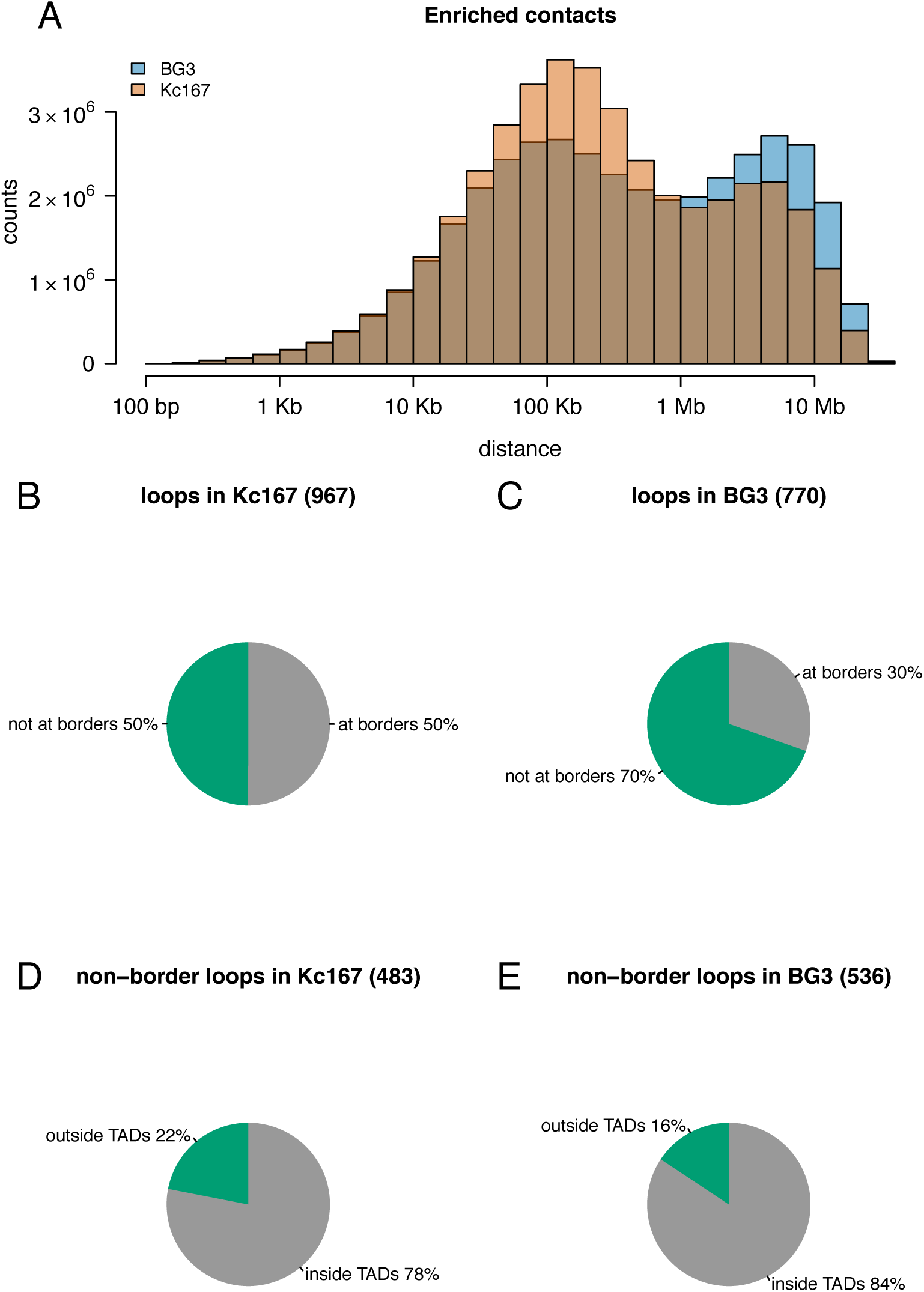
BG3 cells display more long-range interactions compared to Kc167 cells. (A) Histogram showing the distances between two anchors of enriched contacts in the contact matrices (for BG3 and Kc167 cells). Kc167 cells display more contacts for distances between 10 Kb and 1 Mb, while BG3 cells display more contacts for distances over 1 Mb. (B-C) Distribution of chromatin loops at the borders of TADs in Kc167 and BG3 cells showing that Kc167 cells have more loops at TAD boundaries. (D-E) Distribution of non-border chromatin loops within TADs or between TADs in Kc167 and BG3 cells.

We next detected chromatin loops in the two cells types using HICCUPS (Durand et al. 2017) and identified that there are slightly more loops in Kc167 (967) than BG3 (770) cells. Half of the loops in Kc167 cells are at the TAD borders, while only one third of the BG3 loops are located at TAD borders (Figure 6B-C). This suggests that, in Kc167 cells, there are more loop domains (corresponding to a stronger compaction) and in BG3 cells there are more ordinary domains (corresponding to less strict boundaries between domains) (Rao et al. 2014). However, we observed that there are slightly more loops that are inside TADs in BG3 cells compared to Kc167 cells (Figure 6D-E). This suggests that chromatin loops are more present at TAD borders in Kc167 cells, while, in BG3 cells, they are more abundant inside TADs.

Finally, almost half of the loops in both cells have promoters at one anchor, independent of whether the loop is inside TADs or between different TADs, while 10% of the loops have an ncRNA annotated at one anchor (Figure S6). This indicates that active transcription can be involved in maintaining these loops, either by bridging enhancers to promoters or promoters to promoters in order to coordinate gene expression, by the location of genes in transcription factories. In contrast, we found that only a small percentage of loops (10-20%) have an enhancer annotated at one anchor. However, improvements in techniques or analysis tools (Muerdter et al. 2017; Eagen et al. 2017), may likely increase the number of loops that bring together enhancers and promoters.

## DISCUSSION

Recently, several reports came out regarding TADs in *Drosophila melanogaster* (Cubeñas-Potts et al. 2017; Hug et al. 2017; Eagen et al. 2017; Rowley et al. 2017; Ramirez et al. 2017). All these studies generated high resolution Hi-C maps in embryo derived cells or whole embryos. We present here, for the first time an *Insitu* Hi-C map of a differentiated neuronal cell (BG3) in *Drosophila melanogaster* that enabled us to study different aspects of chromatin reorganisation and related features during Drosophila differentiation.

TADs appear similar across various cell types, but there are certain features that dictate cellular identity (which in turn should reflect in the formation of TAD boundaries). Here, with the high-resolution datasets in BG3, we were able to identify TAD borders accurately and compared them with TAD borders in Kc167 cells (embryo derived). Thus, both conserved and new TAD borders gained in BG3 cells were identified (Figure 1E). TADs are regarded as mostly conserved between different cell types (Dixon et al. 2012; Vietri Rudan et al. 2015). We show that, in addition to the similarities, the two different cell types of same organism differ in the nature of TAD borders. Varying transcriptional status and enhancer-promoter interactions across different developmental stages of cells may impact reorganisation of underlying functional elements, which in turn can influence the 3D structure of the genome.

### Role of transcription on TAD border formation

Transcription was considered as a major factor for the basis of TAD formation (Li et al. 2015; Rowley et al. 2017), whereas depletion of expression didn’t seem to affect the TAD formation (Hug et al. 2017). However, presence of concomitant divergent transcription and strong Pol II signal observed at the new BG3 border suggest a strong role for transcription or Pol II to be associated with TAD formation (Figure 2).

Divergent transcription is enriched in mammalian systems and was assumed to be depleted in *Drosophila melanogaster*, but recent reports showed that divergent transcription is more prominent in *D. melanogaster* than previously estimated (Rennie et al. 2017). How would the presence of divergent transcription lead to formation of TADs? One possibility is that divergent transcription produces negative supercoiling in the promoter region of the sense gene, which is maintained during transcription by polymerase, topoisomerase and helicases (Naughton et al. 2013a, 2013b). This is also probably one of the reasons why arresting polymerase by alpha-amanitin did not affect TAD structures (Hug et al. 2017) as alpha-amanitin only blocks the elongation of RNA polymerase II. Additionally, it was shown by numerical simulations using DNA polymers models, that divergent transcription induced supercoiling could explain the self-interacting chromatin domains in *S. pombe* (Benedetti et al. 2017). This hypothesis could be possibly extended to other species like Drosophila and other higher eukaryotes where actual TAD structures may be established and maintained by divergent transcription.

This divergent transcription induced supercoiling was proposed as a mechanism to release paused polymerase in Drosophila (Naughton et al. 2013b). GAF and M1BP seem to be associated with PolII pausing (Li and Gilmour 2013) and, interestingly, we observed strong binding of GAF at conserved TAD borders and an increase of GAF at BG3 specific TAD borders in BG3 cells (Figure S4). Also, it is worthwhile noting that the divergent transcription observed at TAD borders is not only transcription of protein coding genes, but also of ncRNA (Figure S2). This indicates that cell specific ncRNAs could have a role in cell specific TAD border formation or continuance and, consequently, in gene regulation.

### Role of DNA binding proteins at TAD borders

Binding of transcription factors to enhancers activates transcription. Here, we showed that BG3 specific borders display enhancer marks (Figure S5) and are enriched in binding motifs of TFs that are expressed specifically in neuronal cells (Figure 5), thus, supporting that the BG3 specific TAD borders are enriched in BG3 specific enhancers. This indicates that TF binding (including binding of architectural proteins) affects the activity of genes and recruitment of polymerase, consequently, to formation of TAD borders.

Architectural proteins are the other factors, regarded to play a significant role in determining the TAD border formation (Van Bortle et al. 2014). Some of them involved in long-range interactions (such as Chromator and CP190) were enriched at BG3 specific borders validating the presence of distal interactions observed in neuronal cells. Interestingly, in this study, we show that neuronal TAD borders in Drosophila are enriched in CTCF (which commonly occupies mammalian TAD borders and was less pronounced in Drosophila embryonic TAD borders) (Figure 4). The higher occupancy of CTCF in BG3 cells can be explained by higher amounts of CTCF in this cell line (Zabet and Adryan 2015). In combination with Chromator and CP190, they can mediate long-range interactions to meet the transcriptional requirement of the cell during neuronal differentiation. Despite of their presence at the borders, not all architectural proteins were particularly enriched at the neuronal specific borders and therefore may not be essential for the border formation in BG3 cells (Figure S4).

Our data supports the change in reorganisation of the genome observed across different cell stages. TADs are conserved across different organisms and they change during differentiation and development, which probably is the reflection of different transcriptional status, enhancer promoter interactions and differential gene expression. Like in mammalian cells, CTCF might have role in Drosophila that needs to be explored in more detail using different cell type. Furthermore, our results suggest that there is a dynamic change in the enhancer-promoter interaction, especially at BG3 gained borders. This may indicate the role for enhancer promoter interaction as one of the contributing factor for TAD formation. Their organisation across differentiation is also shadowed with cell specific gene expression. Similar observation has been reported in human cells (Bonev et al. 2017; Freire-Pritchett et al. 2017), which suggest that the reorganisation during development is more or less similar and is conserved during evolution. Finally, we hypothesise that active divergent transcription at polymerase paused genes would generate negative supercoiling and in turn may facilitate the formation of compacted domains. The role of the architectural proteins would be to help maintain this negative supercoiling and keep the state of the system stable. However, this still remains an open question and future research might help to elucidate the extended role of each factor in TAD formation.

## METHODS

### Cell Culture

Drosophila BG3 cells were cultured at 25°C in Schneider’s insect medium (Sigma), supplemented with 10% FBS (Labtech), 10 mg/l insulin (Sigma, I9278) and Antibiotic Penstrep.

### Insitu Hi-C protocol

Hi-C libraries were generated from 10 million cells by following the Insitu Hi-C protocol as mentioned in (Rao et al. 2014) with minor modifications. Crosslinked cells were lysed and genome was digested using DpnII (NEB) overnight. The overhangs were filled with Bioton-16-dATP (Jena Bioscience) followed by ligation and de-crosslinking with proteinase K digestion. The sample was further sonicated using Bioruptor. Biotinylated DNA was pulled down using Dynabeads MyOne Streptavidin T1 beads (Life technologies, 65602). Selected biotinylated DNA fragments ranging from 200-500bp were then ligated with illumina adaptors (NEB). The libraries obtained from biological replicates were multiplexed and further sequenced at Edinburgh Genomics (Genepool) using HiSeq4000.

### Hi-C analysis

Each pair of the PE reads were aligned separately to *Drosophila melanogaster* (dm6) genome (Adams et al. 2000; dos Santos et al. 2015) using BWA-mem (Li and Durbin 2010) (with options -t 20 -A1 -B4 -E50 -L0). HiCExplorer was used to build and correct the contact matrices and detect TADs and enriched contacts (Ramirez et al. 2017). The contact matrices were built at 100 Kb bins for plotting Figure 1 and the DpnII restriction sites were used for plotting the rest of the figures. Using a minimum allowed distance between restriction sites of 150 bp and a maximum distance of 800 bp, we obtained a matrix with 217638 bins with a median width of 529 bp. After filtering, the two BG3 replicates had 40 M and 41 M reads, while the Kc167 dataset had 87 M reads (consisting of two pooled replicates). To keep consistency for the comparison, we merged the two BG3 biological replicates, leading to similar number of filtered reads as in the case of Kc167. The matrices were corrected using the thresholds (−1.4 and 5), values were selected from the diagnostic plots (Figure S1A-B). Using the corrected contact matrices, we detected TADs of at least 5 Kb width using a p-value threshold of 0.01 and minimum threshold of the difference between the TAD-separation score of 0.01 (--step 2000, --minBoundaryDistance 5000 --pvalue 0.01 -- delta 0.01). Finally, the enriched contacts were extracted with HiCExplorer using observed/expected ratio method. Same analysis were performed for both the *Insitu* Hi-C datasets of BG3 cells generated in this study and for the *Insitu* Hi-C in Kc167 cells (Cubeñas-Potts et al. 2017). Since, Kc67 cell line is a female derived cell line and BG3 is a male derived cell line, we excluded sex chromosomes from our analysis in order to avoid any biases from dosage compensation (Mukherjee and Beermann 1965; Chiang and Kurnit 2003). The downstream analysis and plots were generated using a custom script in R.

### Motif enrichment analysis

The analysis to identify enriched motifs at TAD borders was performed with R/Bioconductor package PWMEnrich (Stojnic and Diez 2014). First, we created a background model using lognormal method, 200 bp sequence lengths and all Kc167 specific TAD borders (all borders in Kc167 that were further by at least 2 Kb from any TAD border in BG3). Enriched binding motifs that had a p-value lower than 0.001 were selected, which resulted in 66 TFs. Filtering the TF list, we noticed that 3 TFs in the list had 2 motifs so only the motif with the lowest p-value was kept. Finally, using FlyMine (Lyne et al. 2007), we extracted the FlyAtlas expression data of the 63 TFs (Chintapalli et al. 2007).

### Chromatin loops

Chromatin loops were called with HICCUPS tool from Juicer software suite on both Kc167 and BG3 datasets (Durand et al. 2017). In particular, loops were called using 2 Kb resolution, 0.05 FDR, Knight-Ruiz normalisation, a window of 10, peak width of 5, thresholds for merging loops of 0.02,1.5,1.75,2 and distance to merge peaks of 20 Kb (-k KR -r 2000 –f 0.05 -p 5 -i 10 -t 0.02,1.5,1.75,2 -d 20000); for details on the parameters see (Durand et al. 2017).

### Datasets

Note that due to the similarities in chromatin nature and transcriptional profiles of the Kc167 and S2 cell types (Cubeñas-Potts et al. 2017), depending on data availability, we used one of the two as the embryonic cell line, when comparing to neuronal cell line (BG3). To keep consistency with our TAD annotation, where files had coordinates in other release versions of *Drosophila melanogaster* genome, the coordinates were lifted to dm6.

#### ChIP-chip

We used the following ChIP-chip datasets generated and preprocessed (M values smoothed over 500 bp) by modEncode consortium (modENCODE Consortium et al. 2010; Kharchenko et al. 2010; Riddle et al. 2011; Schwartz et al. 2012): PolII (GSE20832, GSE20806), BEAF- 32 (GSE20811, GSE20760), CP190 (GSE20814, GSE20766), Chromator (GSE20761, GSE20763), CTCF (GSE20767, GSE32818), GAF (GSE23466, GSE32822), Su(Hw) (GSE20833, GSE51964), H3K27ac (GSE20778, GSE20779), H3K27me3 (GSE20780, GSE45083), H3K4me1 (GSE23468, GSE45085), H3K4me3 (GSE20839, GSE45088).

#### Transcription

The mRNA abundance in the two cell lines was downloaded from (Lee et al. 2014), who preprocessed the original modEncode datasets (Cherbas et al. 2011). To obtain the strand specific expression, we mapped the genes on the corresponding strand, using the FlyBase annotation (dos Santos et al. 2015). For nascent RNA transcription, we used the preprocessed GRO-seq in S2 cells (GSM577244) from (Core et al. 2012) and the preprocessed 3’NT-seq in BG3 cells (GSE100545) from (Pherson et al. 2017).

#### Other datasets

We also used gene and ncRNA annotations for Figure S2 from Flybase (dos Santos et al. 2015), STARR-seq annotation of enhancers in BG3 and S2 cells from (Arnold et al. 2013; Yanez-Cuna et al. 2014) and preprocessed DNaseI-seq profiles from modEncode (Kharchenko et al. 2010).

## Data availability

*Insitu* Hi-C dataset was deposited on GEO.

## Acknowledgements

This work was supported by Wellcome Trust grant 202012/Z/16/Z. We would like to thank Professor Sarah Bray and her lab, especially Zoe Pillidge, Matthew Jones and Dr Lenka Skalska for useful discussions and support with BG3 cell line culture. Also we would like to thank Dr Rob White for useful discussion and comments on the project. The analysis was performed on the HPC at University of Essex and we would like to thank Stuart Newman for his support on using the cluster. Also we would like to thank Professor Christopher Reynolds for providing access to his GPU machine to run HICCUPS. We thank Ilya M Flyamer for helping us with Hi-C protocol.

## Author Contributions

Conception and design of the experiments: KC and NRZ. Performed the experiments: KC. Analysis of the data: NRZ. Written the paper: KC and NRZ.

